# Length-dependent translation efficiency of ER-destined proteins

**DOI:** 10.1101/2023.03.16.532890

**Authors:** Hana Sahinbegovic, Alexander Vdovin, Renata Snaurova, Michal Durech, Jakub Nezval, Jiri Sobotka, Roman Hajek, Tomas Jelinek, Michal Simicek

## Abstract

Gene expression resulting in the generation of new proteins is a fundamental process critical for every living organism. Particularly in eukaryotic cells, complex organization of the cell body requires fine-tuning of every step prior to de novo protein synthesis. To ensure proper localization, certain mRNAs possess unique signal sequence, which destinies the translation apparatus to the specific organelle. Here we focus on the mechanisms governing the translation of signal sequence-bearing mRNAs, which encode proteins targeted to the endoplasmic reticulum (ER). The binding of a signal-recognition particle (SRP) to the translation machinery halts protein synthesis until the mRNA-ribosome complex reaches ER membrane. The commonly accepted model suggests that mRNA containing the ER signal peptide continuously repeats the cycle of SRP binding followed by association and dissociation with ER. In contrast with the current view, we show that the long mRNAs remain on the ER while being translated. On the other hand, due to a low ribosome occupancy, the short mRNAs continue the cycle always facing the translation pause. Ultimately, this leads to a significant drop in the translation efficiency of small, ER-targeted proteins. The proposed mechanism advances our understanding of selective protein synthesis in eukaryotic cells and provides new avenues to enhance protein production in biotechnological settings.

## Introduction

Translation is a fundamental and tightly-regulated process used by every cell to produce proteins by decoding the genetic information from the form of mRNA transcript into an amino acid sequence in the polypeptide. A single transcript can be read by multiple ribosomes at the same time forming a complex known as polysome (Munro et al. 1964). Once engaged, polysome will translate the mRNAs and would not readily exchange it for newly added mRNA until the translation of the initial mRNA is completed (Thompson and Gilbert 2017). Transcripts associated with multiple ribosomes suggest robust translation. On the other hand, mRNAs with low or no ribosome associations are expected to be translated poorly or remain untranslated (Panda et al. 2017). Additionally, the presence of rare codons, complex secondary structures, premature stop codons or missense sequences can result in ribosome collisions and stalling events during the translation elongation phase (Arpat et al. 2020). Subsequently, inhibition of translation triggers ribosome quality control and mRNA surveillance pathways that ultimately lead to the dissociation of ribosomes and degradation of nascent peptide (Shoemaker and Green 2012).

The endoplasmic reticulum (ER) plays a vital role in the correct assembly of proteins destined for membrane integration or secretion. The ER-directed mRNA encompasses about 30-40% of all eukaryotic transcripts (Pyhtila et al. 2008). The majority of these mRNAs does not bind to ER itself and their association with ER is enabled by the binding interactions of the ribosome and translational product. The most well-described mechanism for ER localization of mRNA relies on the presence of a specific stretch of amino acids known as signal peptide (SP) which upon translation is recognized by a signal recognition particle (SRP) (Lingappa and Blobel 1980; Blobel 2000).

During translation, polypeptides are translocated into ER lumen which significantly differs from the cytosol with respect to ion concentration and redox conditions. Importantly, ER lumen promotes a variety of post-translational modifications and chaperone-facilitated folding and is thus crucial for the proper assembly and function of most of the membrane and secretory proteins (Ellgaard et al. 2016). Proteins detected as terminally misfolded are destined for degradation by the cytosolic ubiquitin-proteasome system (Nandi et al. 2006). Moreover, protein quality control in ER stops protein synthesis while simultaneously expanding the capacity for folding and degradation in response to stress (Sitia and Braakman 2003). The implementation of strict control measures in the ER guarantees the capture of potentially harmful proteins (McCaffrey and Braakman 2016).

In this study, we uncover a novel phenomenon of the length-dependent translation efficiency for the ER-destined SP-containing mRNAs. We found that protein synthesis into ER from short mRNAs is significantly less efficient compared to longer mRNAs. The apparent translation deficiency for shorter, ER-localized mRNAs is not affected components of the ribosome stalling pathway. We also show that cells tend to upregulate the expression of shorter transcripts which is however not fully compensating for a drop in protein synthesis. Further, our work suggests that mRNAs encoding short proteins possessing ER-localisation SP are less associated with ER membrane and excluded from polysome fractions. Altogether, we propose a model that suggests the translation mechanism for short and long ER-destined transcripts.

## Results

### Single IgL domain decreases translation efficiency similar to poly-A sequence

To date, ribosome stalling was associated mainly with mRNA pathologies such as the presence of premature stop codons or complex secondary structures. Several studies have also suggested the need for translation slowdown in the interdomain regions to allow correct protein folding. This might increase the chances of unwanted ribosome crushes. To further understand, whether multidomain proteins might be prone to ribosome collisions, we used a well-established fluorescent reporter system. Initially, we replaced the poly-A stalling sequence with one, two and four Ig domains of the Igλ light chain (IgL) (Fig 1A). Surprisingly, a single IgL domain interrupted translation to a similar extent as the poly-A cassette. On the other hand, two and four IgL domains had no effect on the translation efficiency (Fig 1B). To reveal, whether the observed translation inhibition is related to protein fold, we tested the effect of structurally related Ig-like domain from Filamin A (FLNA) (Fig 1C). Through similar size and 3D arrangement, the presence of neither one nor multiple FLNA Ig-like repeats did not replicate the translation interruption caused by a single IgL domain (Fig 1D).

**Fig 1.**
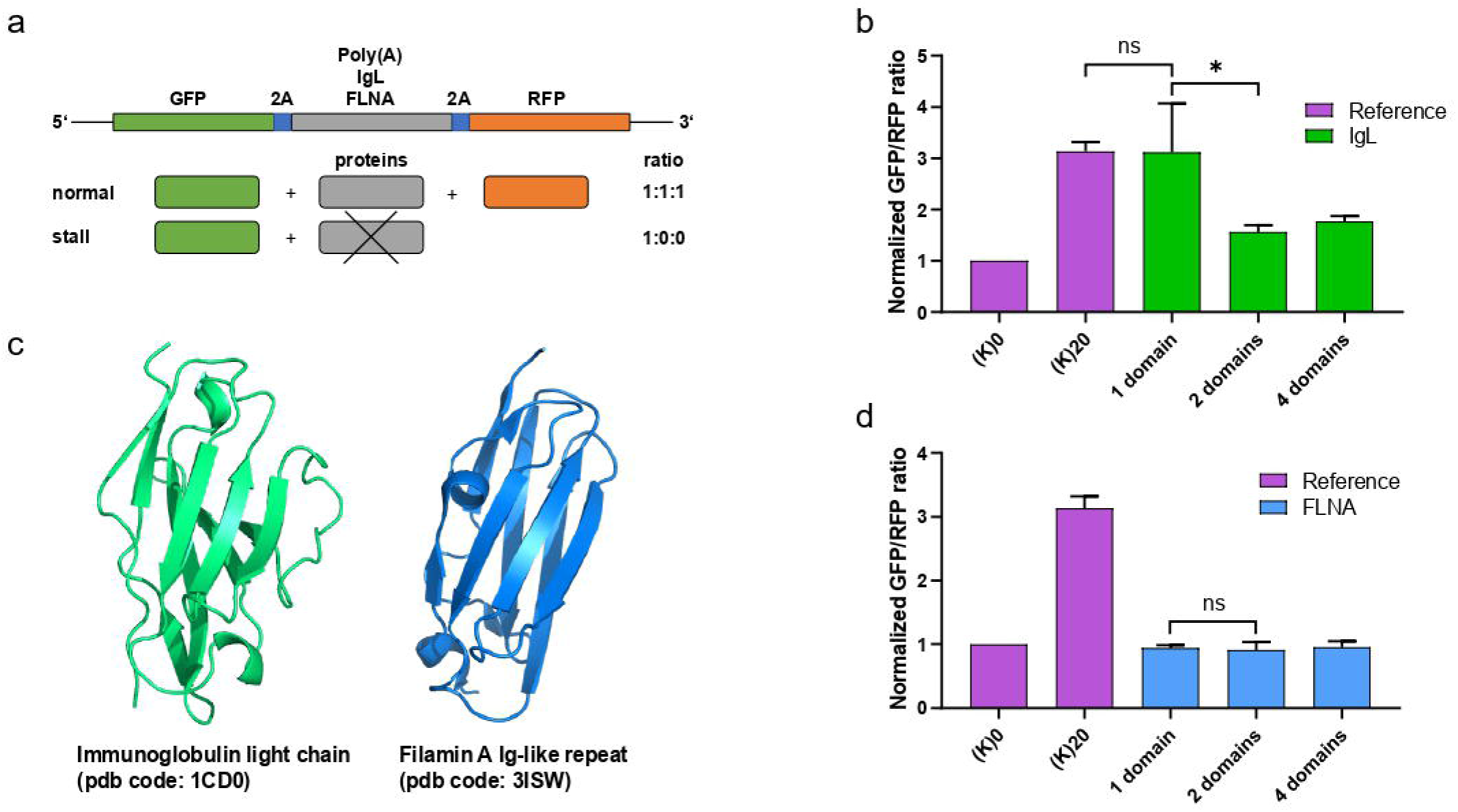
Single Ig domain abrogates translation. **(A)** Schematic representation of the translation efficiency reporter system with various stalling cassettes. **(B)** Normalized GFP:RFP fluorescence ratio of the translation efficiency reporter containing the control (K)0 and stalling (K)20 cassettes (Reference, purple) and repeats of 1, 2, 4 IgLλ domains (IgL, green). Significance was compared using the two-tailed Student’s t test, * = P, 0.05. Error bars represent the mean ±SD. **(C)** X-ray structures of IgLλ (green, pdb code: 1CD0) and Filamin A (blue, pdb code: 3ISW). **(D)** Normalized GFP:RFP fluorescence ratio of the translation efficiency reporter containing the control and stalling cassettes (K)0 and (K)20 (Reference, purple) and repeats of 1, 2, 4 FLNA Ig-like domains (FLNA, blue). Significance was compared using the two-tailed Student’s t test, * = P, 0.05, ns = non-significant. Error bars represent the mean ±SD.

### Translation of IgL does not induce ribosome stalling

To examine if the observed phenomenon is mechanistically related to the canonical ribosome stalling pathway, we overexpressed the E3-ligase ZNF598 which can inhibit translation through ubiquitination of small ribosome subunit RPS10. Interestingly, ZNF598 failed to block translation in our single IgL domain reporter system (Fig 2A). Complementary, we analyzed the ubiquitination of endogenous RPS10, which is one of the initial events triggered by stalled ribosomes. As expected, RPS10 ubiquitination increased upon expression of the reporter with the poly-A sequence (K)20 but not a single IgL domain. These results suggest that inhibition of translation mediated by one IgL domain is likely not connected to ribosome stalling. Given no effect of sequentially and structurally highly related FLNA Ig-like repeat, we speculated that subcellular localization of particular mRNA might play a role in translation efficiency. Therefore, we deleted the ER destination signal peptide from IgL (ΔSP IgL) and fused the same signal sequence to the Ig-like domain of FLNA (SP FLNA). In accordance with our hypothesis, the swap of targeting SP rescued translation block mediated by a single IgL domain while inducing partial translation inhibition in presence of one Ig-like domain from FLNA. Together, our data indicate that the translation efficiency of short mRNAs containing ER-targeting SP might be significantly diminished.

**Fig 2.**
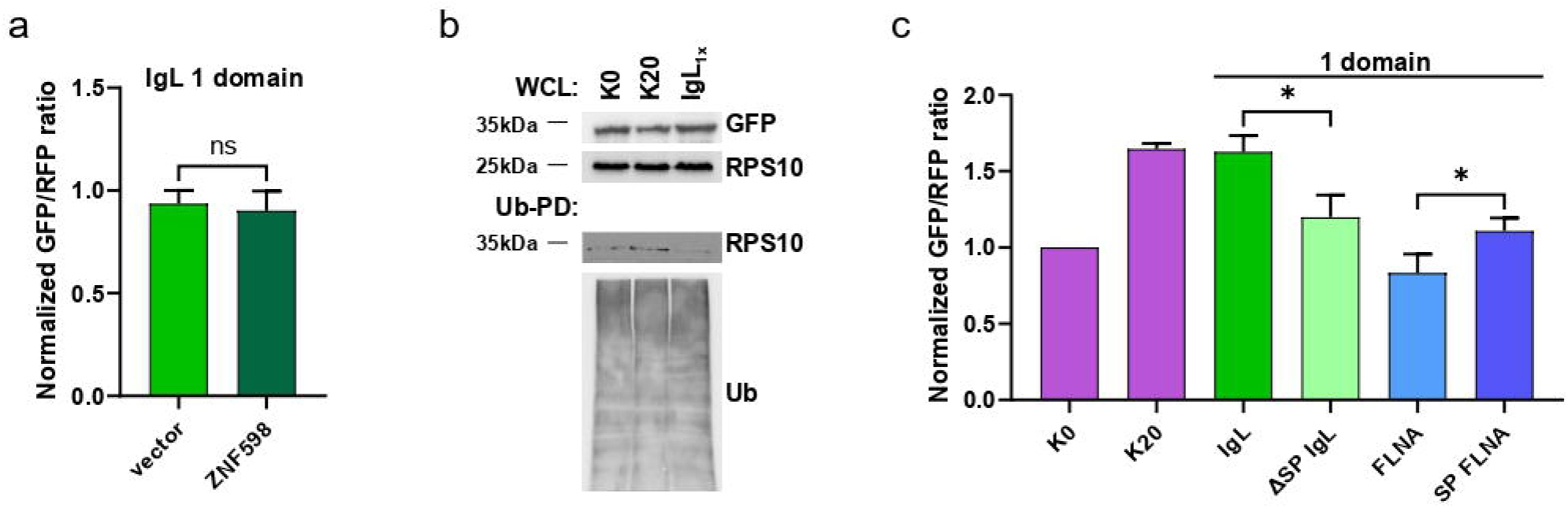
Evaluation of ribosome stalling and ER SP on translation efficiency of short mRNA. **(A)** Normalized GFP:RFP fluorescence ratio of the translation efficiency reporter containing a single repeat of IgLλ domain (IgL) together with co-expression of empty vector (light green) or ZNF598 (dark green). Significance was compared using the two-tailed Student’s t test, ns = non-significant. Error bars represent the mean ±SD. **(B)** Ubiquitin pull-down (Ub-PD) and whole cell lystate (WCL) from cell expressing the translation efficiency reporter containing the control (K)0, stalling cassettes (K)20 (Reference) and single repeat of IgLλ domain (IgL_1x_). **(C)** Normalized GFP:RFP fluorescence ratio of the translation efficiency reporter containing the control (K)0 and stalling (K)20 cassettes (purple), single repeat of IgL (green) and FLNA Ig-like domains with deletion (ΔSP IgL) and insertion of SP (SP FLNA). Significance was compared using the two-tailed Student’s t test, * = P, 0.05, ns = non-significant. Error bars represent the mean ±SD.

### The amount of short ER-destined proteins does not correlate with the respective mRNAs

In order to determine whether the destination of translation to ER might globally affect protein synthesis, we quantified the abundance of various mRNAs based on the presence of SP. Analysis of the expression dataset (Cunningham et al. 2022) revealed a clear trend of enhanced transcription towards shorter mRNAs. This tendency was particularly prominent in mRNAs encoding proteins with ER-targeting SP (Fig 3A). The absolute quantity of proteins mostly correlated with the relative amount of their respective mRNAs (Fig 3B). The exception were proteins whose corresponding mRNA open reading frame (ORF) was shorter than 400 nucleotides (nt). In this group, the number of synthesized proteins did not correlate with the relative amount of matching mRNAs. This suggests that short mRNAs (ORF < 400nt) containing ER-targeting SP might indeed be translated less efficiently compared to longer mRNAs.

**Fig 3.**
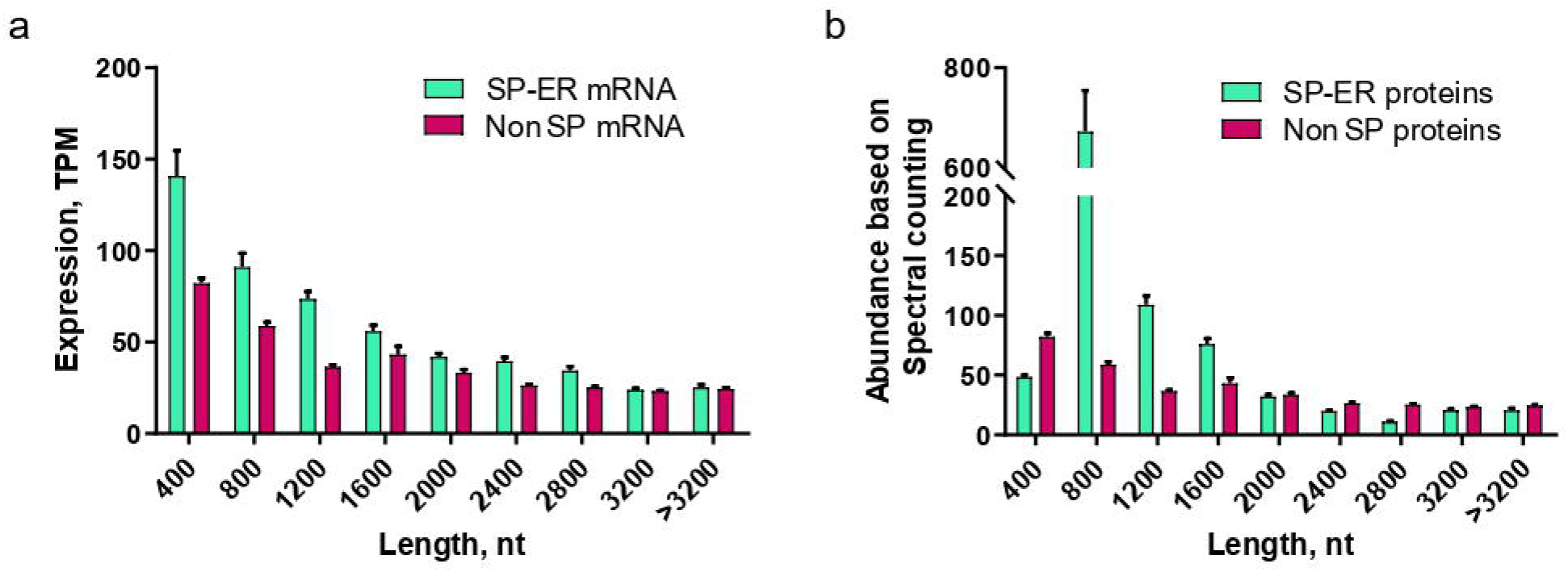
Analysis of transcript and protein abundance. **(A)** Average expression of human genes (transcripts per million reads) from 32 tissues in 122 individuals. Genes are divided into groups based on the length of the open reading frame. **(B)** Average abundance of human proteins (parts per million), whole organism (integrated). Proteins are divided into groups based on the length of the relative coding sequence.

### Length-dependent association of mRNA with intracellular membranes

Given the apparent difference in translation efficiency of short *vs* long ER-destined proteins, we speculated that the length of mRNA might affect the association of translation machinery with ER membrane. To this end, we performed cell fractionation to separate cytosolic and membrane-bound mRNAs (Fig. 4A). Initially, we used cells with expression of single or tandem IgL domains. Upon quantification based on qPCR analysis of IgL mRNA, we found that the mRNA encoding two IgL domains was more abundant in the membrane fraction compared to mRNA with single IgL domain ORF (Fig. 4B). To validate this observation, we monitored the presence of endogenous mRNAs with ER-targeting SP and monitor their presence in cytosolic and membrane pools. The detected mRNAs were divided into two groups: short (>400nt) and long (<400nt). We found that the membrane to cytosol ratio was higher for longer mRNAs compared to short mRNAs (Fig. 4C). These results indicate that the length of mRNAs with ER-destination SP might define the interval of active translation on ER membrane.

**Fig 4.**
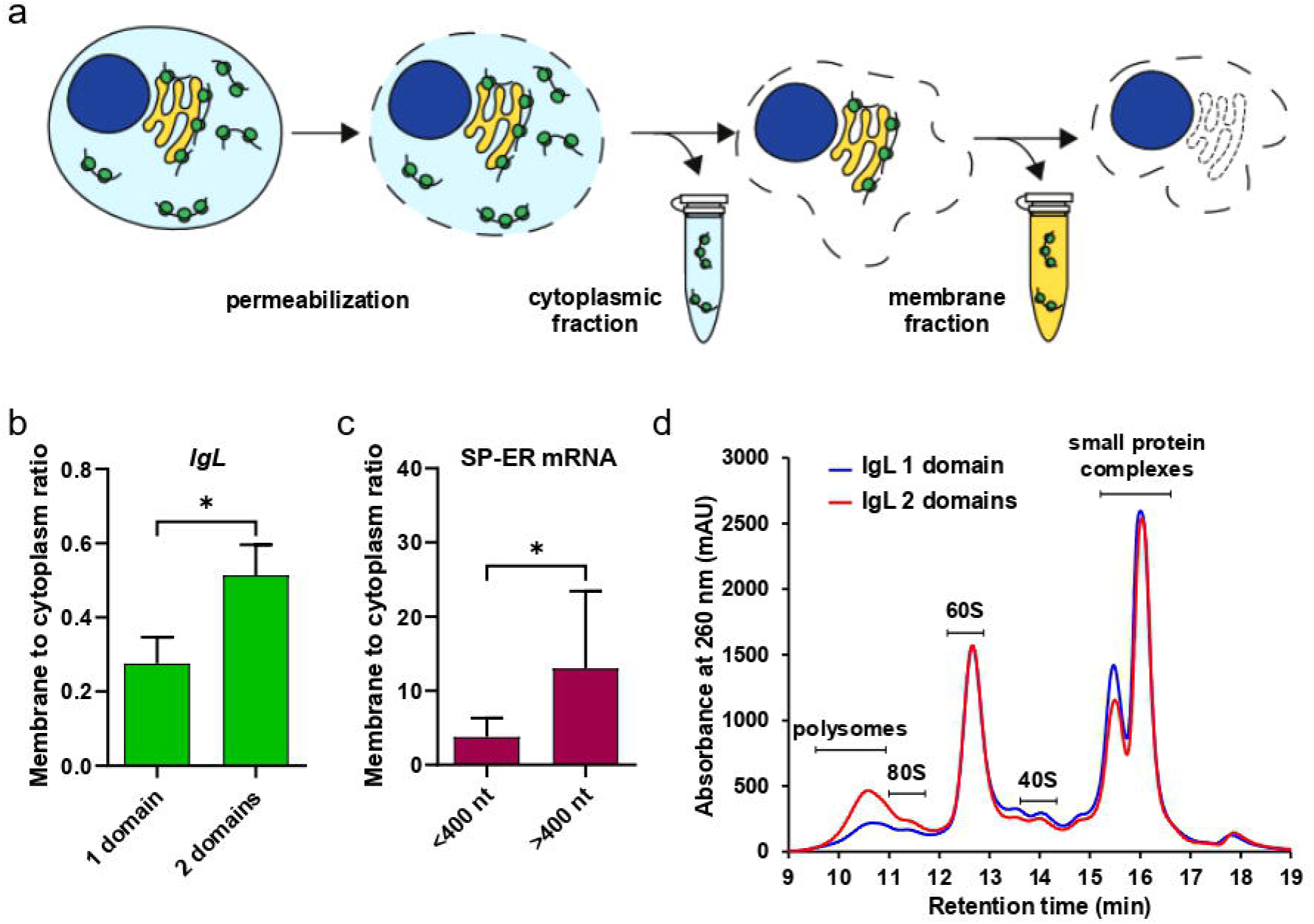
Membrane localisation of short *vs* long mRNAs. **(A)** Schematic representation of the fractionation assay. **(B)** Relative ratio of free (cytosolic) and membrane-bound (ER-associated) mRNA encoding 1 and 2 IgLλ domains. Significance was compared using the two-tailed Student’s t test, * = P, 0.05, ns = non-significant. Error bars represent the mean ±SD. **(C)** Relative ratio of free (cytosolic) and membrane-bound (ER-associated) endogenous mRNA encoding short (<400 nt, n= 8 genes) and long (>400 nt, n = 9 genes) proteins containing ER-destination SP. Significance was compared using the two-tailed Student’s t test, * = P, 0.05, ns = non-significant. Error bars represent the mean ±SD. **(D)** Polysome analysis using Ribo Mega-SEC. Cell lysates from HEK293 cells transfected plasmid encoding 1 and 2 IgLλ domains were separated using Agilent Bio SEC-5 2000 Å column. The retention time is indicated on the x axis and the UV absorbancde at 260 nm is shown on the y axis.

The 400 nt is approximately the average distance between two elongating ribosomes on the eukaryotic mRNA. Thus, we hypothesized that short mRNA (<400nt) might be occupied with only one ribosome while longer mRNA (>400nt) contains multiple active translation complexes. To test this assumption, we analyzed polysome profiles from cells with overexpression of single and tandem IgL domains (Fig 4D). As expected, the cells with abundant mRNA encoding two IgL domains resemble significantly enriched polysomal pool compare to cells expressing single IgL ORF.

## Discussion

Translation of transmembrane, ER luminal or secreted proteins starts in cytosol, where the general translation factors and ribosomal subunits bind to mRNA. After the synthesis of SP the nascent polypeptide and ribosome-mRNA complex are recognized by SRP, which triggers a temporary pause in the proteosynthesis. The SRP-dependent elongation arrest acts until the proper localization of the translating ribosome on the ER membrane is ensured (Walter and Blobel 1981). Moreover, translation arrest is crucial taken that SRP cannot target proteins when the nascent polypeptide exceeds a critical length (Flanagan et al. 2003). SRP-bound translation machinery engages with SRP receptor on the ER-membrane where the nascent polypeptide chain is positioned into translocon allowing active insertion of newly formed protein into the ER lumen. Meanwhile, SRP and SRP receptor are released from the translation complex (Akopian et al. 2013).

Location of majority of the SP-containing mRNAs to ER membrane is fully dependent on the assembled translational complex that associates with SRP (Walter and Blobel 1981). If the SP is present on the protein N-terminus, it is usually chopped off by ER proteases soon after it appears inside ER. Once the protein synthesis is completed, the ribosome dissociates from mRNA allowing new translational cycle to begin. Although the process is not well understood yet, accumulating evidence also points to the existence of SP-independent localisation of the translation apparatus to the ER membrane (Pyhtila et al. 2008; Reid and Nicchitta 2015).

The translation efficiency can be regulated at various levels including by the number of elongating ribosomes on particular mRNA (Li et al. 2014). The minimal distance between actively translating ribosomes in eukaryotic cells is assumed to be around 400 nt (Yan et al. 2016). Thus, the short mRNAs (ORF<400nt) encoding proteins directed to ER can be occupied only by a single elongating ribosome at the time. Therefore, disassembly of the translation apparatus would likely lead to the dissociation of the short mRNA from the ER membrane to the cytosol. Re-initiation of the translation cycle will result in the synthesis of the SP sequence and binding of SRP followed by a translation pause until the entire complex reaches the ER membrane.

On the other hand, longer SP-containing mRNAs (ORF>400nt) can be processed by multiple active ribosomes forming polysome assembly, where the translation complexes associate with the ER translocon at various elongation state. The continuous binding of translating ribosomes might allow for persisting localization of the long mRNAs on the ER membrane. Based on our data we hypothesize that due to the close proximity of the SRP-bound ribosomes with the respective translocons, the temporary translation pause caused by SRP biding is minimized. This ultimately results in increased efficiency of proteosynthesis from the longer mRNAs compared to the short mRNAs (Fig 5).

**Fig 5.**
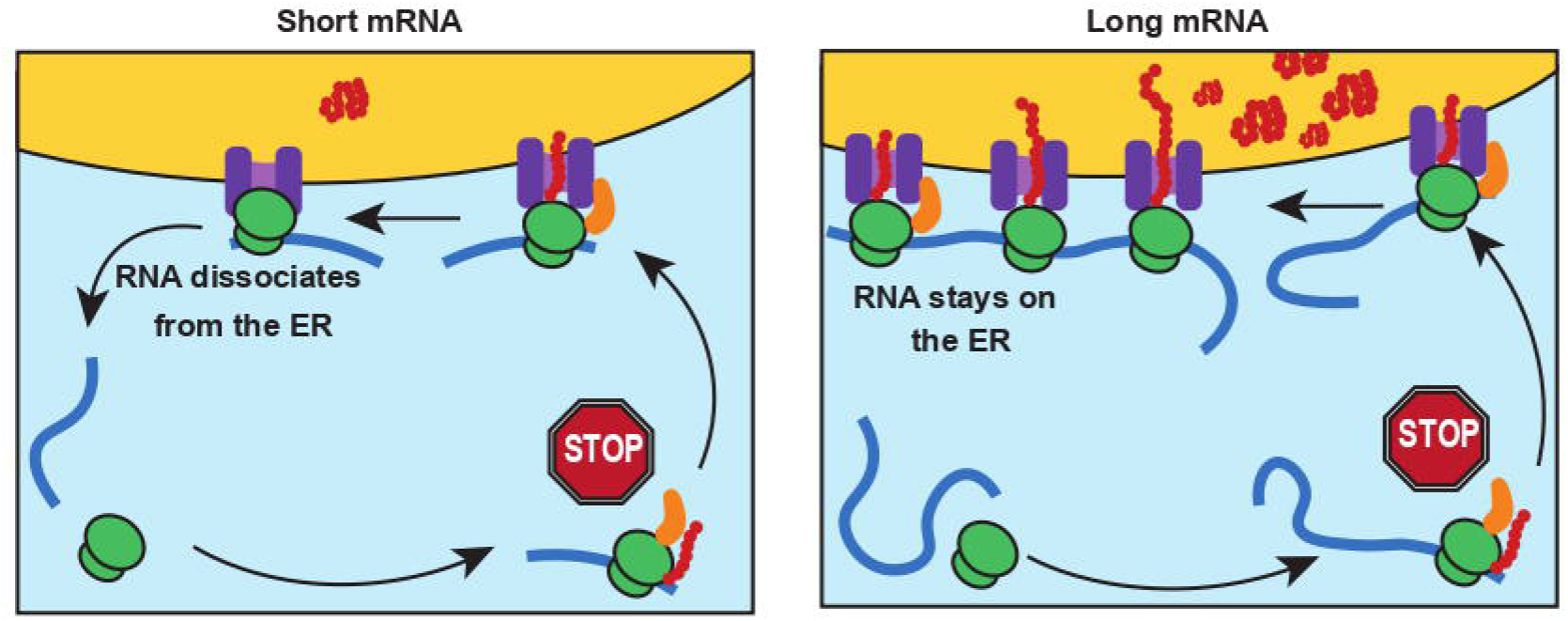
Schematic model of length-dependent translation efficiency of the ER-destined proteins. The proposed model of SP-dependent localisation of short *vs* long ER-targeted mRNA.

In accordance with our model, it was previously shown that transcripts with higher ribosome densities have a higher rate of translation. However, mRNAs with short ORF are some of the most highly translated in the cell (Thompson et al. 2016). The apparent discrepancy is usually explained by the closed-loop model of ribosome-mRNA translation complex. In this scenario, the eukaryotic mRNA is recruited to the ribosome via recognition of the 5’m7GpppN cap by translation initiation factor eIF4F, which can interact with the 3’poly(A)-binding proteins (PABP) bringing the two ends of mRNA in close proximity (Tomek and Wollenhaupt 2012).

Moreover, shorter mRNAs experience higher ribosome re-initiation rates due to higher enrichment with closed-loop factors, thus forming more closed-loop complexes and enabling the ribosomes to reinitiate the translation on the same transcript multiple times (Rogers et al. 2017). It is therefore possible that short mRNAs contain a common attribute such as *cis*-acting motif that allows hasty and preferential recruitment of closed-loop factors. Such motif has however never been identified. Diffusion was also suggested, as it would allow the end collision of shorter molecules, giving the closed loop more opportunities to form (Guo et al. 2014).

Another possibility is that ribosomes sense the length of ORF and thus promote the interaction with closed-loop factors (Thompson and Gilbert 2017). This length-dependent mechanism could have several advantages. Genes with the housekeeping role tend to be encoded in short ORFs (Eisenberg and Levanon 2003; Thompson and Gilbert 2017) and their expression has to be constant. Additionally, a closed loop would allow the cell to conserve resources in poor growth conditions (Panda et al. 2017).

This model, however, does not include specific subcellular localization of particular mRNA, which might interfere with the interaction of translation initiation factors and PABP proteins. Specifically, SRP-dependent delivery and subsequent association of translation apparatus with proteins on the ER membrane might be challenging for the stability of the closed-loop structure. Therefore, to fully delineate the mechanism behind the lower translation efficiency of short ER SP-containing mRNAs, additional studies are needed. Nevertheless, our findings provide potential new avenues to enhance the synthesis of small secreted proteins in eukaryotic expression systems.

## Materials and methods

### Cell lines, culture conditions, and transfections

HEK293 cells were maintained in Dulbecco’s Modified Eagle’s Medium (DMEM; high glucose, pyruvate, and L-glutamine) containing 10% fetal bovine serum (FBS) and 1% penicillin/streptomycin and maintained in a 5% CO_2_ humidified incubator. Cells were transiently transfected with the respective plasmids using polyethyleneimine (PEI; 1 mg/mL) and serum-free media (Opti-MEM), mixed, and incubated 15 min at RT when cells were at 60% confluence 24 h after seeding.

### Quantitative real-time PCR

QIAzol Lysis Reagent (Qiagen) was used to extract total RNA. The RNA samples’ quality (purity and integrity) was determined utilizing the Agilent 2100 Bioanalyser with the RNA 600 NanoLabChip reagent set (Agilent Technologies). The RNA was quantified by Qubit fluorometry (Thermo Scientific). The RevertAid First Strand cDNA Synthesis Kit (Thermo Scientific) was used for Complementary DNA (cDNA) synthesis according to the manufacturer’s instructions. Quantitative PCR was conducted using PowerUpTMSYBRTM Green Master Mix (Applied Biosystems) on StrepOnePlus real-time PCR system (Applied Biosystems). Relative mRNA expression was calculated by 2-DDCt method and normalized to GAPDH transcripts.

### Flow cytometry analysis

Translation efficiency analysis employing flow cytometry was performed using the stalling reporter system as described previously (Juszkiewicz and Hegde 2017). Briefly, 10^6^ HEK293 cells transfected with a dual-fluorescent reporter containing (K)0 or (K^AAA^)20, or reporter with one, two or four domains of IgL, or one, two or four domains of FLNA, or one domain of IgL with ZNF598, were trypsinized after 72 h transfection using TrypleX, sedimented (600 g for 3 min), washed in PBS, sedimented again and resuspended in PBS containing SYTOX blue and analyzed using a Beckman Coulter’s CytoFLEX S instrument. Approximately 10^4^ events were collected in SYTOX negative gate for each sample, geometric mean statistics were used to analyze MFI in the FlowJo software.

### SDS-PAGE and Western blotting

The cell lysates were resolved by SDS PAGE and transferred to the polyvinylidene difluoride (PVDF) membrane. Membranes were blocked in 5% (wt/vol) nonfat milk (Roth) in PBS-T (phosphate buffer saline, 0.05% Tween 20) and incubated with the appropriate primary antibodies overnight at 4°C in 1% (wt/vol) BSA/PBS-T. Primary antibodies used at indicated dilutions include anti-GFP (MMS-118P, Biolegend, 1:2000), anti-RPS10 (sc-515655, SantaCruz, 1:1,000) and anti-Ub (3936, Cell Signaling Technology, 1:1000). Membranes were subsequently washed with PBS-T and incubated with horseradish peroxidase (HRP)-conjugated secondary antibody goat anti-mouse-IgG (115-035-146, Jackson ImmunoResearch, 1:2,000) for 1 h at room temperature. Signal detection was performed using ECL (Thermo Fisher Scientific) and ChemiDoc MP System (Bio-Rad). Protein bands analysis by densitometry utilized ImageJ v1.49 (National Institutes of Health).

### Preparation of cell lysates for Ribo Mega-SEC

A total of 2*10^7^ HEK293 transfected with reporter system were treated with 50 µg/mL cycloheximide to maintain polysome stability for 5 min under 37°C and 5% CO_2_ before harvest. Cells were then washed with ice-cold PBS containing 50 µg/mL cycloheximide, lysed by vortexing for 10 s in 400 µL of polysome extraction buffer (20 mM HEPES-NaOH, pH 7.4, 130 mM NaCl, 10 mM MgCl_2_, 1% CHAPS,2.5 mM DTT, 50 µg/mL cycloheximide, 20 U RNase inhibitor murine, New England BioLabs, EDTA-free protease inhibitor cocktail, Thermo Scientific) and incubated for 30 min on ice. Insoluble material was pelleted by centrifugation at 17,000 g for 10 min at 4°C. The supernatant was filtered through 0.45 µm Ultrafree-MC HV centrifugal filter units by 12,000 g for 2 min, and the total RNA amount in the filtrate was quantified by Qubit fluorometry (Thermo Scientific).

### Polysome separation by Ribo Mega-SEC

Agilent Bio SEC-5 column (2000 Å, 7.8 × 300 mm, 5 µm particles) together with Agilent Bio SEC-5 guard column (2000 Å, 7.8 × 50 mm, 5 µm particles) was connected to the Agilent 1200 HPLC system (Agilent Technologies) and equilibrated with 2 column volumes (CV) of filtered SEC buffer (20 mM HEPES-NaOH, pH 7.4, 100 mM NaCl, 10 mM MgCl_2_, 0.3% CHAPS, 2.5 mM DTT). 100 µL of 10 mg/mL of filtered bovine serum albumin (BSA) solution diluted by SEC buffer was injected once to block the sites for nonspecific interactions. After monitoring the column condition by injecting 25 µL of GeneRuler 1 kb DNA Ladder (Thermo Scientific), 100 µL of cell lysate containing 120 µg of RNA was loaded on the column. All column conditioning and separation were done at 12°C. The chromatogram was monitored by measuring UV absorbance at 215, 260, and 280 nm by the diode array detector. The flow rate was 0.8 mL/min.

### Fractionation by sequential detergent extraction

Fractionation of the HEK293 cells was done according to the protocol by Jagannathan et al (Jagannathan et al. 2011). Briefly, HEK293 cells at 80-90% confluency were treated with ice-cold PBS containing 50 µg/mL cyclohexamide for 10 min, followed by incubation with 1 mL of ice-cold permeabilization buffer for 5 min. Soluble material was collected as the cytosolic fraction. Cells were then washed with ice-cold wash buffer and incubated with 1 mL of ice-cold lysis buffer for 5 min. The soluble fraction was collected as a membrane fraction. Both cytosolic and membrane fractions were centrifuged at 7,500 × g for 10 min and clear supernatants were transferred to new tubes.

### Databases

Data on mRNAs with ER-signal peptide was obtained from BioMart (Cunningham et al. 2022). Data for protein abundance was taken from PAXdb: Protein Abundance Database (Wang et al. 2015).

### Statistical analysis

The statistical significance of differences between various groups was calculated with the two-tailed unpaired t-test. Error bars represent the standard deviation of the mean (SD). Statistical analyses, unless otherwise indicated, were performed using GraphPad Prism 5.

## Acknowledgments

This work was financially supported by the Grant Agency of the Czech Republic (GA CR 21-21413S) and by the Cell Coolab Ostrava - Research and Development Center for Cell Therapy in Hematology and Oncology (No. CZ.02.1.01/0.0/0.0/17_049/0008440) finances provided from ERDF.

